# Life-history traits as predictors of expected genetic contributions 15 years later in a cooperatively breeding bird

**DOI:** 10.64898/2025.12.19.695490

**Authors:** EA Young, E Chesterton, MA Versteegh, M van der Velde, SF Worsley, T Burke, J Komdeur, DS Richardson, HL Dugdale

**Author notes:** Corresponding authors –. Equal first authorship (EA Young, E Chesterton).

## Abstract

An individual’s life history plays an important role in how successful individuals are in passing on their genes to future generations. However, exactly which life-history traits best approximate an individual’s long-term genetic contributions remains poorly studied, especially in cooperative breeders. Here, we use a genetic pedigree and long-term data of the closed population of the Seychelles warbler (*Acrocephalus sechellensis*) to calculate the expected individual genetic contributions (IGC) after 15 years (or ∼3 generations) across 11 cohorts. Using a Bayesian analysis we then quantified and compared how well six important life-history traits predict IGC in each sex of this cooperatively breeding species. The life-history traits compared were: acquisition of a dominant breeding position, age at first breeding attempt, tenure as a dominant breeder, lifespan, and LRS (measured as both the number of independent [surviving 3 months+] and recruited [surviving 1 year+] offspring produced over an individual’s lifetime). An individual’s lifetime reproductive success (LRS) was the strongest predictor of IGC and this predictive power did not differ across the different measurements used. However, this predictive power did differ between sexes, explaining ∼61-62% of IGC variation in males versus ∼47-49% in females owing to the larger variation among males in their annual reproductive success. Across other life-history traits the predictive power of IGC was similar across both sexes but differed between life-history traits considerably (2-37%). Comparing these predictive powers suggested that lifespan and the duration of a dominant breeding position are far more important in shaping long-term genetic contributions than if and when in their life they acquired dominance. Overall, these findings give insight into the individual and population-level processes influencing future gene pools and illustrate how life-history evolution is shaped by sex-specific reproductive patterns and cooperative breeding dynamics.

## INTRODUCTION

Fitness, a concept with profound consequences in evolutionary biology, is broadly concerned with the relative ability of a genotype or phenotype to persist and proliferate through generations due to natural selection (Grafen, 1988; Williams, 1966). To measure how selection acts on a particular phenotype or genotype, researchers often measure how these units of selection are associated with components of an individual’s life-history, such as their survival or reproductive performance (e.g., Grant, 1985). However, these fitness proxies provide only a snapshot of evolutionary processes that act over much longer timescales, and we have a poor understanding of how well they predict individual long-term contributions to the gene pool (Brommer et al., 2004; Reid et al., 2019).

Recent studies have used long-term pedigree data on wild animal populations to examine how well different fitness proxies predict an individual’s expected genetic contributions [IGC; (Alif et al., 2022; Chen et al., 2019; Reid et al., 2019; Young et al., 2023) - the proportion of the gene pool that an individual has contributed through direct descent after several generations, based upon the pedigree data (Hunter et al., 2019)]. IGC will partly reflect an individual’s fitness - individuals surviving longer and reproducing more should have higher IGC. However, IGC will also reflect demographic (e.g., population increases/decreases or rates of immigration/emigration) and stochastic processes [e.g., genetic drift (Reid et al., 2019)]. Thus, comparing how well different life-history traits predict IGC can help to: (1) elucidate the role of natural selection versus other evolutionary forces (i.e., gene flow and genetic drift) in shaping an individual’s long-term contributions to the gene pool, and (2) evaluate how well different life-history traits capture selection.

An individual’s lifetime reproductive success (LRS) – the total number of offspring an individual produces during its lifetime (Clutton-Brock, 1988) – is the life-history trait generally agreed to be the gold standard for measuring the direction and strength of natural selection (Crow, 1958; Lande & Arnold, 1983). Consequently, LRS is the most widely used fitness proxy (Kingsolver et al., 2001). However, when measuring LRS, researchers face decisions such as when to count, or census, offspring. For example, in avian species, LRS has been quantified as the number of eggs (e.g., Mumme, 1992), hatchlings (e.g., Alif et al., 2022), nestlings (e.g., Safina & Burger, 1983), fledglings (e.g., Gates & Gysel, 1978), independent offspring (e.g., Raj Pant et al., 2022) or offspring recruited into the adult or breeding population (e.g., Meierhofer et al., 1999) produced by an individual over the course of its life. Censusing offspring earlier in life comes with the advantages of estimating fitness closer to its theoretical zygote-to-zygote definition (Evans & Postma, 2024), reduces underestimating viability selection (Grafen, 1988; Hadfield, 2008), and avoids conflating the fitness of parents and offspring (Wolf & Wade, 2001). However, if producing a higher quantity of offspring comes with significant costs to the quality of offspring (Lack, 1947), then censoring offspring later in life may be necessary to avoid biasing estimates of fitness. Indeed, previous studies have typically shown that calculating LRS using offspring censused at a later life-history stage increases its predictive ability for IGC (Alif et al., 2022; Chen et al., 2019; Reid et al., 2019; but see Young et al., 2023). However, this may be because later measures of LRS can better account for random variation in the survival of offspring to these later stages, rather than strong quantity–quality trade-offs *per se* (Young et al., 2023). Moreover, these trends have only been identified within three different wild animal populations, and so the ability of different measures of LRS to predict IGC compared to other life-history traits, in different species, remains poorly understood (Alif et al., 2022).

Studies have also explored how well lifespan as a life-history trait predicts IGC, but results have varied across species. For example, lifespan was a stronger predictor of IGC in song sparrows [*Melospiza melodia*, explaining 25-29% of variation in IGC (Reid et al., 2019)] than in pre-industrial humans, where lifespan explained only ∼13% of the variation in IGC (Young et al., 2023). This is logical, considering that humans tend to stop reproducing well before the end of their lifespan (Hawkes et al., 1998), whereas, in birds, individuals are capable of reproducing for a much greater proportion of their lives, and so lifespan plays a more important role in shaping IGC, especially when variance in annual reproductive success is low. However, as lifespan does not account for an individual’s reproductive output, it is still expected to be a worse predictor of IGC than LRS [(Reid et al., 2019), see also Supplementary Table 1 (Young et al., 2023)]. Lifespan will also become a worse predictor in the face of stronger reproductive senescence – whereby older individuals have lower annual reproductive success (Dugdale et al., 2011) and quality of offspring (Lansing, 1947; Monaghan et al., 2020; Sparks, Spurgin, et al., 2022) – and reproductive–survival trade-offs (Stearns, 1989; Williams, 1966). However, to what extent lifespan predicts IGC, relative to LRS, deserves investigation across more species and across different life histories.

Cooperative breeders have a unique life history - where subordinate helpers care for the offspring of others - and deserve special attention when examining how well different life-history traits predict IGC. However, to our knowledge, only one study has examined how well different life-history traits predict IGC in a cooperatively breeding species (Chen et al., 2019), focusing only on measures of LRS and the numbers of grandoffspring and great-grandoffspring, without comparing these measures to other life-history traits. Other life-history traits may also place constraints on fitness in cooperative breeding species. For example, many cooperative breeders are socially hierarchical (Komdeur et al., 2016), with dominant individuals being responsible for most of the population’s reproductive output, and subordinate individuals helping to raise the offspring of dominant breeders to gain indirect benefits such as an increased chance of gaining a future dominant breeding position (Chesterton et al., 2024). Thus, LRS may be constrained by: (1) whether individuals attain a dominant breeding position; (2) the age at which individuals gain a dominant breeding position; and (3) how long this tenure lasts. As no studies have examined how these different life-history traits predict IGC relative to LRS in cooperative breeding species, we have a limited understanding of how life-history patterns shape the ability of different fitness proxies to predict IGC and their interplay with other evolutionary processes.

To explore how well life-history traits predict IGC in a cooperatively breeding bird, we use the long-term Seychelles warbler (*Acrocephalus sechellensis*) dataset. The closed (Komdeur et al., 2004) population of this species on Cousin Island has a high resighting rate (98% for adults, Brouwer et al., 2010) and assigned genetic parentage (Hadfield et al., 2006; Richardson et al., 2001; Sparks, Hammers, et al., 2022), meaning that the pedigree and individual estimates of survival and reproductive success are highly accurate, which is rare in natural populations. We estimate how much variance in future genetic contributions can be explained by single-generation life-history traits. To do this, we compare how well different life-history traits predict IGC within the population after 15 years (∼3 generations) across 680 individuals. The six life-history traits used were: (A) acquisition of a dominant breeding position, (B) age at first breeding attempt, (C) tenure as a dominant breeder, (D) lifespan, and (E) LRS (measured as both the number of independent [surviving 3 months+] and (F) recruited [surviving 1 year+] offspring produced over an individual’s lifetime, Table 1). We did this separately for both sexes, modelling IGC within a zero-inflated Bayesian model framework. This allowed us to determine how each life-history trait affected whether an individual’s lineage survived after 15 years (i.e., lineage extinction probability), the IGC of individuals whose lineages did not go extinct, and whether the predictive power of life-history traits was different for each trait or across males and females.

**Table 1.**
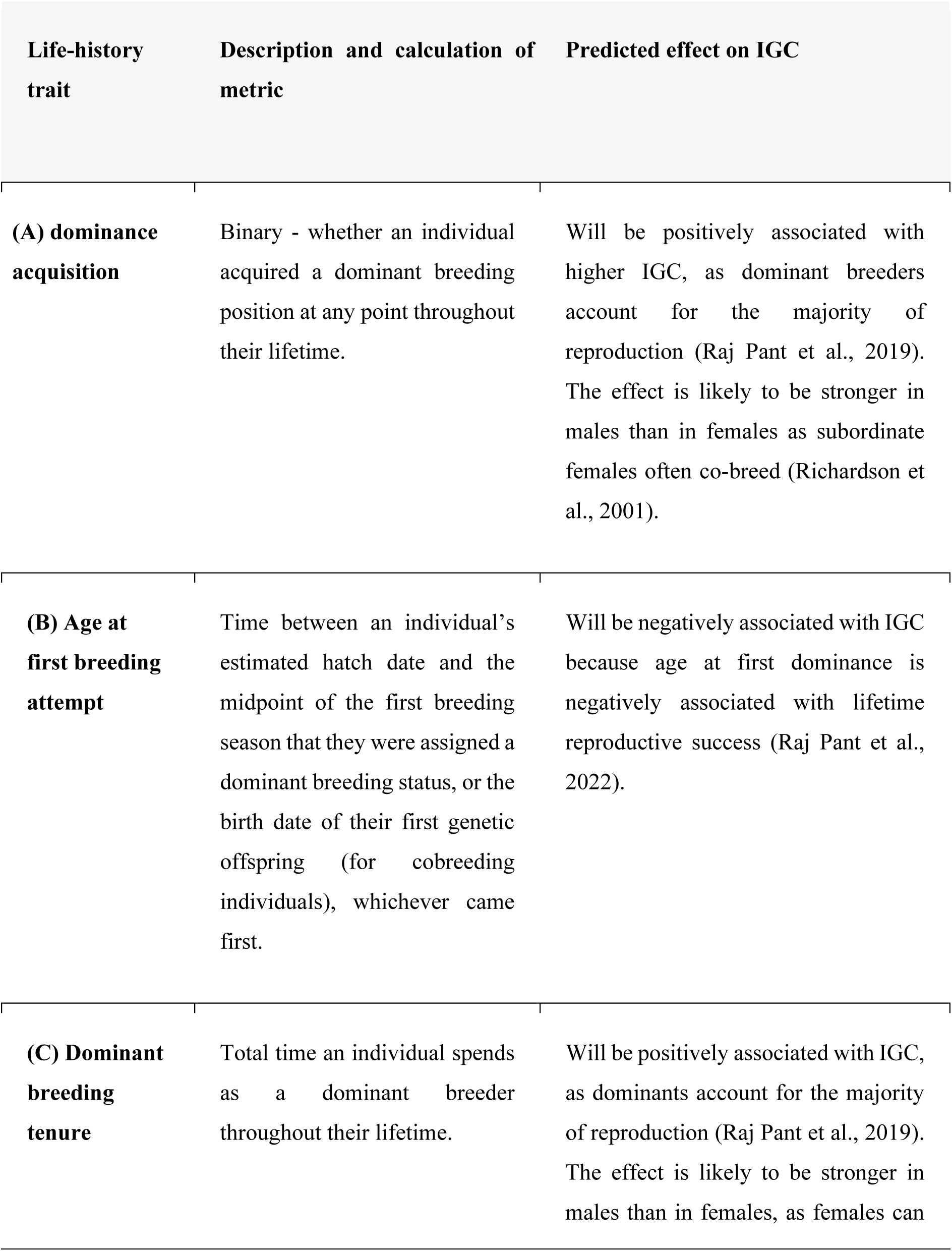
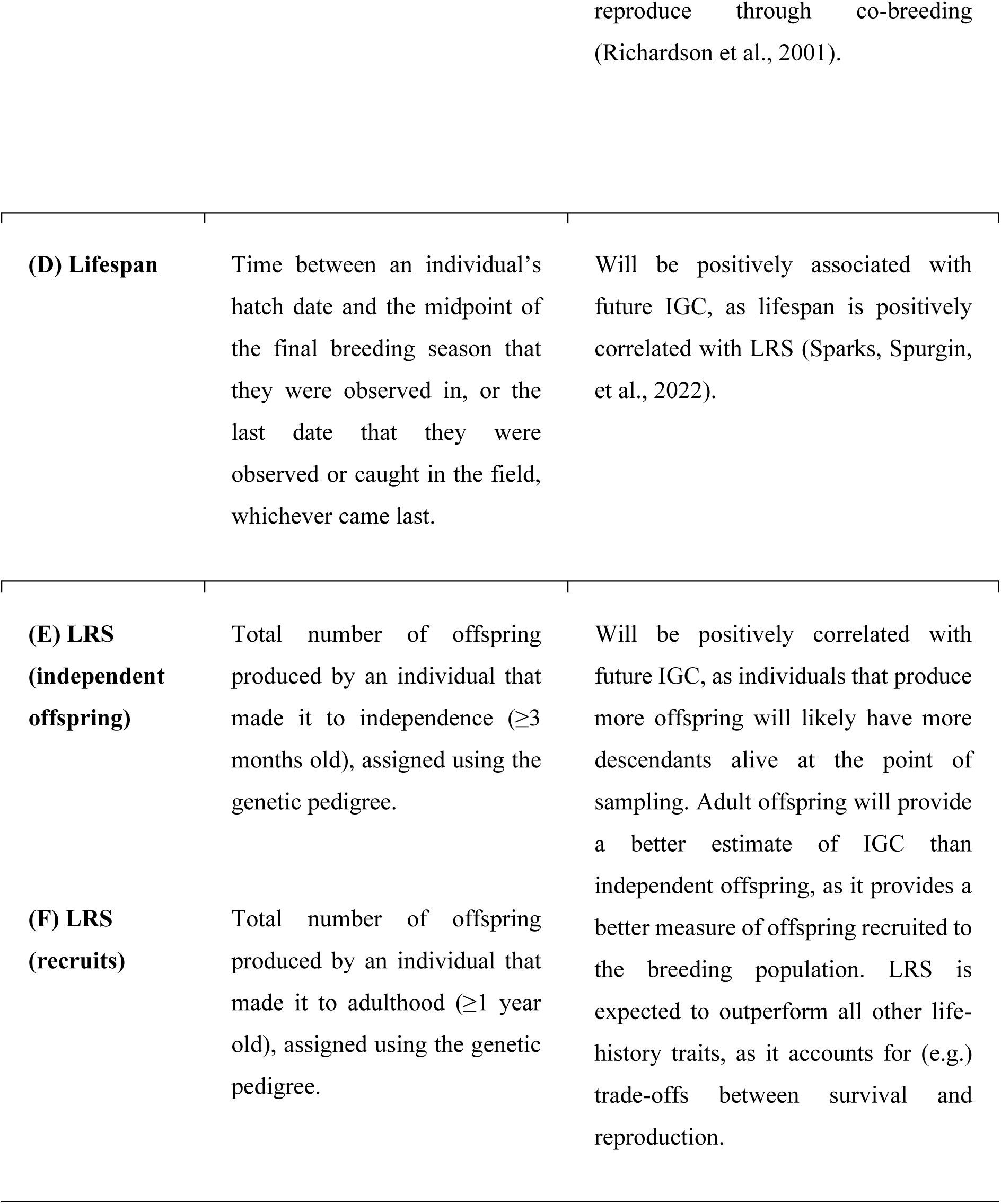
Descriptions of single-generation life-history traits, how they were calculated, and their predicted effect on future individual genetic contribution (IGC) to a population 15 years in the future in the Seychelles warbler.

## METHODS

### Study system and data collection

The Seychelles warbler is a facultative cooperatively breeding passerine endemic to the Seychelles archipelago (Komdeur et al., 2016). The closed population on Cousin Island (0.29 km^2^; 4°20′S, 55°40′E) consists of approximately 320 adults which remained stable at carrying capacity from 1997-2022, and has been studied since 1985, with intensive monitoring from 1997 onwards (Davies et al., 2021; Richardson et al., 2002). Each year, the population is usually monitored over two breeding seasons: the major (June–September) and minor (January–March) breeding seasons, respectively (Richardson et al., 2002). There are normally ∼110 Seychelles warbler territories on Cousin (Komdeur, 2003), each of which is occupied by a breeding group consisting of a dominant breeding pair and 0-5 subordinates, some of which help to raise the offspring (Komdeur, 1992; Richardson et al., 2002). The statuses of individuals are determined through behavioural observations during the breeding seasons. Dominant breeders are determined by observations of mate guarding and contact calls (Richardson et al., 2002). Subordinates are defined as all other sexually mature adults within a territory.

During the breeding seasons, as many Seychelles warblers as possible are captured in the nest as nestlings or in mist nets. The majority of birds (96%; Richardson et al., 2001; Sparks, Spurgin, et al., 2022) have been ringed with a unique combination of three ultraviolet-resistant colour rings and a British Trust for Ornithology metal ring. Blood samples (*ca.* 25 μl) are taken using brachial venepuncture and stored in 100% ethanol at room temperature. Molecular sexing is performed with 1–3 markers, and 30 microsatellites are used to genotype individuals and determine genetic parentage (Sparks, Spurgin, et al., 2022). The R package MasterBayes 2.52 (Hadfield et al., 2006) was used to assign parentage using methods detailed in Sparks, Spurgin, et al., (2022). Of the 2,282 birds hatched between 1992 and up to and including the major breeding season in 2022, 88% of fathers and 85% of mothers have been assigned at ≥80% accuracy. Importantly, the microsatellites used for parentage assignment are under neutral selection (Supplementary methods; Fig S1). From these parentage data, a pedigree spanning 12 generations was created (Raj Pant et al., 2022; Sparks, Spurgin, et al., 2022), from which accurate information about reproductive success could be extracted.

### Single-generation life-history traits

The fitness component estimates from the Seychelles warbler are unusually accurate for a wild population. This is because the whole island population is intensively monitored, the resighting probability is high (0.92 ± 0.02 for <2-year-olds and 0.98 ± 0.01 for older birds; Brouwer et al., 2010), and inter-island dispersal is rare (<0.1% of all individuals studied, Komdeur et al., 2004). Therefore, if an individual is not seen for ≥2 consecutive field seasons, we can confidently assume that it is dead (Brouwer et al., 2006). The separation of dispersal and death allows lifetime fitness estimates, such as lifespan and LRS, to be accurately calculated.

For each individual, we considered how acquiring dominance, the age at first breeding attempt, dominance tenure, lifespan, and LRS (measured as both the number of independent [≥3 months] and adult [≥1 year] offspring) predicted the estimated IGC to the population 15 years (∼3 generations) later (see Table 1 for full definitions of life-history traits, and their predicted relationship with IGC). The number of independent and adult offspring were both considered to assess whether censoring offspring at later stages provided a better estimate of IGC by accounting for juvenile mortality, as observed in three other systems (Alif et al., 2022; Reid et al., 2019; Young et al., 2023). Our core dataset included focal individuals hatched over 11 years (1997–2007) during both the major and minor breeding seasons. Only individuals hatched from 1997 onwards were included, as this is when intensive monitoring began, with a great majority of individuals being genotyped and having parentage assigned (Sparks, Spurgin, et al., 2022). Removing individuals hatched before 1997 reduced the likelihood of LRS being underestimated. Because Seychelles warblers have overlapping generations, analyses focused on yearly cohorts, rather than discrete generations. Only individuals that survived to independence (≥3 months) were included, of which all could be caught, to remove potential bias in the dataset; nests located high in the canopy are hard to find and access, so individuals hatched in high nests might not be sampled until after fledging, and catching effort has varied across years, such that offspring have been first caught at different ages (Raj Pant et al., 2022). Our final (largest) dataset consisted of 337 females and 343 males.

### Estimation of individual genetic contributions

Individual genetic contributions (IGC) were calculated using a pedigree-based method following Hunter et al., (2019). Parents - at least autosomally - each contribute 50% of their genes to offspring. This method extends these expectations to estimate the expected genetic contribution to more distant direct descendants: e.g., on average, 25% for grand-offspring, 12.5% for great-grand-offspring, etc. These genetic contributions are then totalled up for all living descendants present in the population at a specific point in time for each focal individual. For example, an individual that has three grand-offspring alive in the population at the time point studied would have an expected genetic contribution of 0.75 (0.25 x 3), plus 1 if it was itself also still alive within the population. This value can then be divided by the total population size at that point to allow comparisons between cohorts and time periods. We calculated IGC 15 years post-hatching for each focal individual, enabling us to estimate the IGC for 680 focal individuals across 11 cohorts (1997–2007; Fig. S2), approximately 3 generations into the future. Generation time was calculated as the sex-specific mean parental age at hatching in our dataset (females: 4.7 years, males: 5.1 years, following Young et al., 2023).

### Statistical analyses

To examine how well each life-history trait predicted the IGC 15 years post-hatching, we ran generalised linear mixed models (GLMMs) with zero-inflated beta distributions using the log and logit-link functions. Using zero-inflated beta models allowed us to examine whether the life-history traits predicted the probability of lineage extinction (i.e., an individual having no IGC after 15 years), and - if the lineage persisted - the value of the IGC. For both the zero-inflated and beta model components, we used the Probability of Direction (*pd*) parameter to examine whether life-history traits predicted IGC. Specifically, we considered pd values <0.95 as not significant, 0.95-0.975 as marginally significant, and >0.975 as significant (Makowski et al., 2019). We performed a Benjamin-Hochberg adjustment for multiple testing on pd values (Verhoeven et al., 2005), per sex and distribution (i.e. 6 tests). Models were run for each life-history trait and sex separately. To evaluate whether different life-history traits in each sex better predicted IGC, we estimated the Bayesian R^2^ from our models and examined whether the 95% credible intervals for the differences in R^2^ (ΔR^2^) between the models for each sex overlapped zero (Gelman et al., 2019). Cohort was included as a random effect to control for temporal variation in mean IGC.

All models were run with brms 2.22.0 (Bürkner, 2018) using the Markov Chain Monte Carlo sampler in RStan 2.32.6.6 (Stan Development Team, 2020) in R 4.3.3 (R Core Team, 2020). 2020). Across all models, four chains were run for 9,000 iterations, sampling every 15 iterations after burning the first 2,000 samples. We used default uninformative priors: fixed effects, flat; random effects, Student’s t-distribution. Convergence was visually inspected using caterpillar plots and by confirming that all Rhat parameters were equal to 1. We also checked model fit using the brms *pp_check* function. Marginal effects plots were created using *conditional_effects()* from brms. We calculated Bayesian R^2^, and the difference of the R^2^ values between the life-history traits with *bayes_r2()* from the brms package. AI was not used in these analyses.

## RESULTS

Of 680 focal warblers born 1997-2007, 64.7% of females (*n* = 218/337) and 69.1% of males (*n* = 237/343) had zero IGC in the population after 15 years (i.e., their lineages went extinct, Figure S2). For both females and males, acquiring dominance, longer dominance tenures, longer lifespans, and higher LRS (of both independent and recruited offspring) reduced the likelihood of lineage extinction (*pd* > 0.975) whereas age at first breeding attempt did not affect the likelihood of lineage extinction (*pd* < 0.975, Table 2; Figure 1). For individuals whose lineages did not go extinct, longer dominance tenures, longer lifespans, and higher LRS (both independent and recruited offspring) also increased IGC in both sexes (*pd* > 0.975, Table 2; Figure 2). Age at first breeding attempt also did not affect the IGC of individuals whose lineages did not go extinct in either sex (pd < 0.975), (*pd* > 0.975, Table 2; Figure 2A, Figure B).

**Figure 1.**
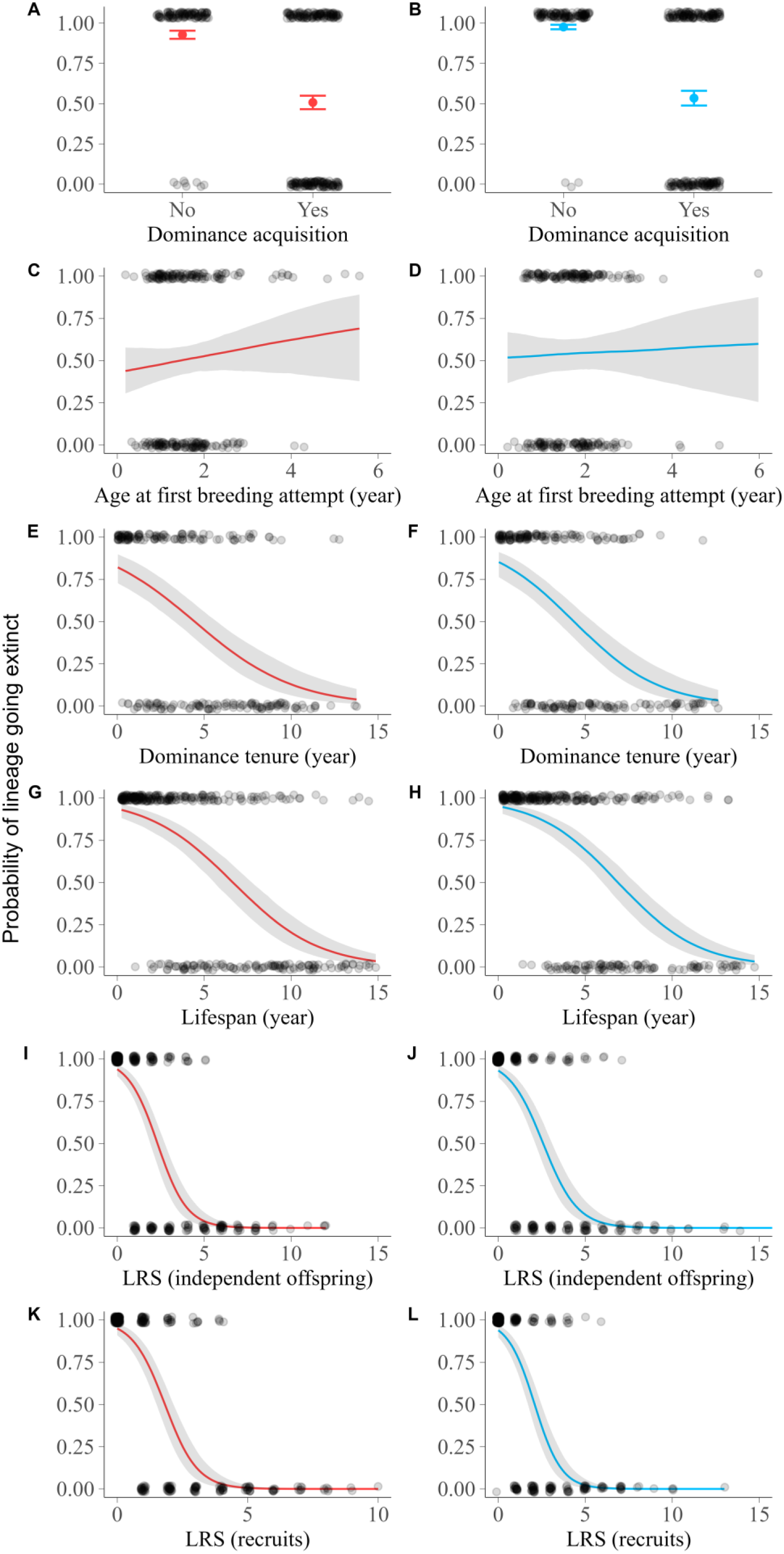
The relationship between the probability of the lineage going extinct 15 years in the future for female (red) and male (blue) Seychelles warblers and (A, B) dominance acquisition (females and males, respectively), (C, D) age at first breeding attempt, (E, F) dominance tenure, (G, H) lifespan (years)), (I, J) LRS (independent offspring, ≥3 months old), and (K, L) LRS (recruits, ≥1 year old). Probabilities are based on whether the focal individual was still alive and the estimated proportion of genetic information that the individual shared with all of their living, direct descendants in the sample cohort 15 years in the future. Individual values (dots) are presented slightly jittered to aid visualisation alongside the fitted regression line. The grey shading around the fitted regression line shows the corresponding 95% credible interval.

**Figure 2.**
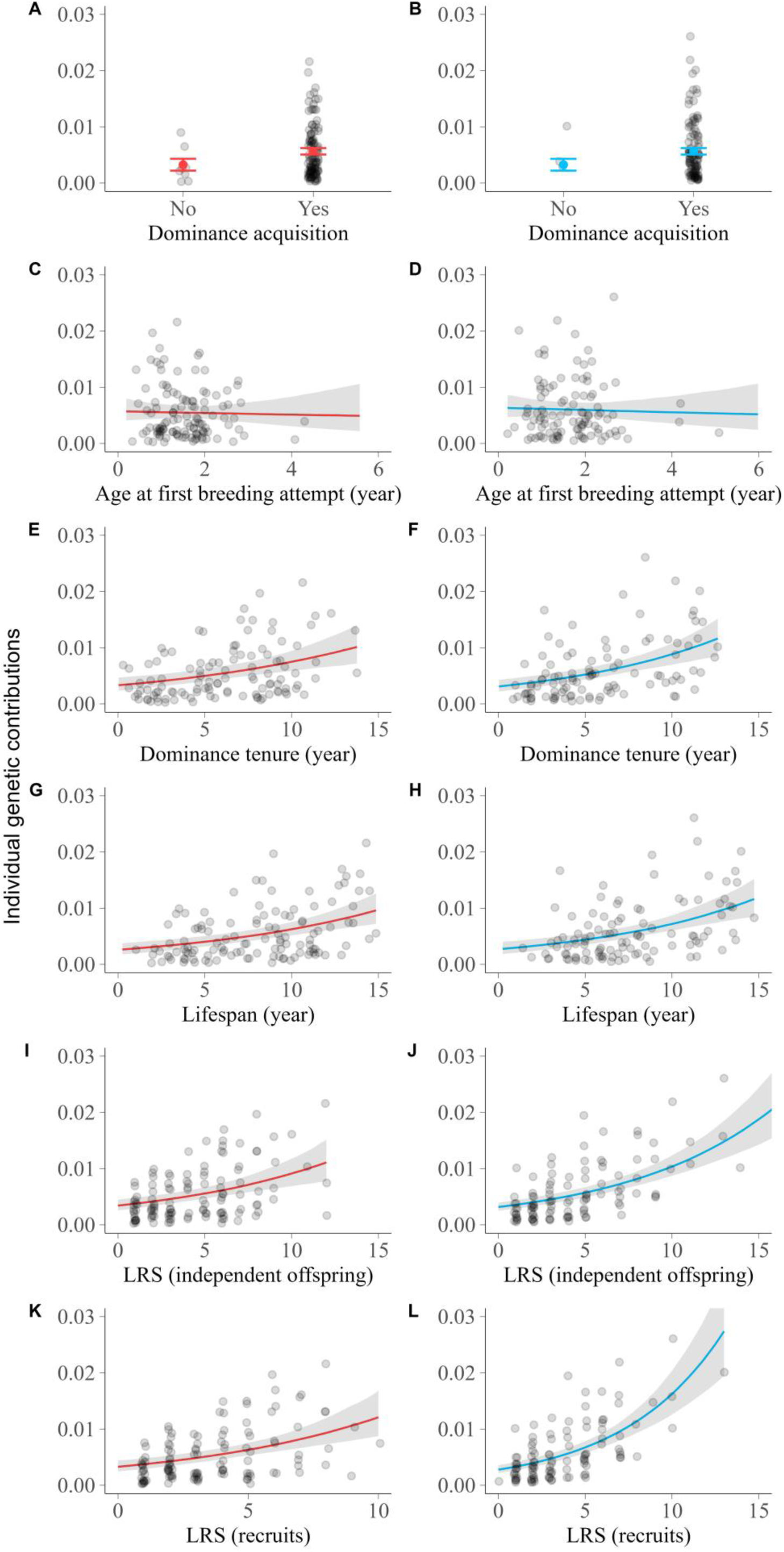
The relationship between the estimated non-zero individual genetic contribution (IGC) to a population 15 years in the future for Seychelles warblers and (A,B) dominance acquisition (female, red, and male, blue, respectively), (C, D) age at first breeding attempt, (E, F) dominance tenure, (G, H) lifespan (years), (I, J) LRS (independent offspring, ≥3 months old), and (K, L) LRS (recruits, ≥1 year old). Estimated IGC was calculated by considering the estimated proportion of genetic information that an individual shares with all of their living, direct descendants (i.e., the summed *r* coefficient), divided by the total number of individuals alive in the sample cohort 15 years in the future. Individual values (dots) are presented, along with the fitted regression line. The grey shading around the fitted regression line shows the corresponding 95% credible interval.

**Table 2.**
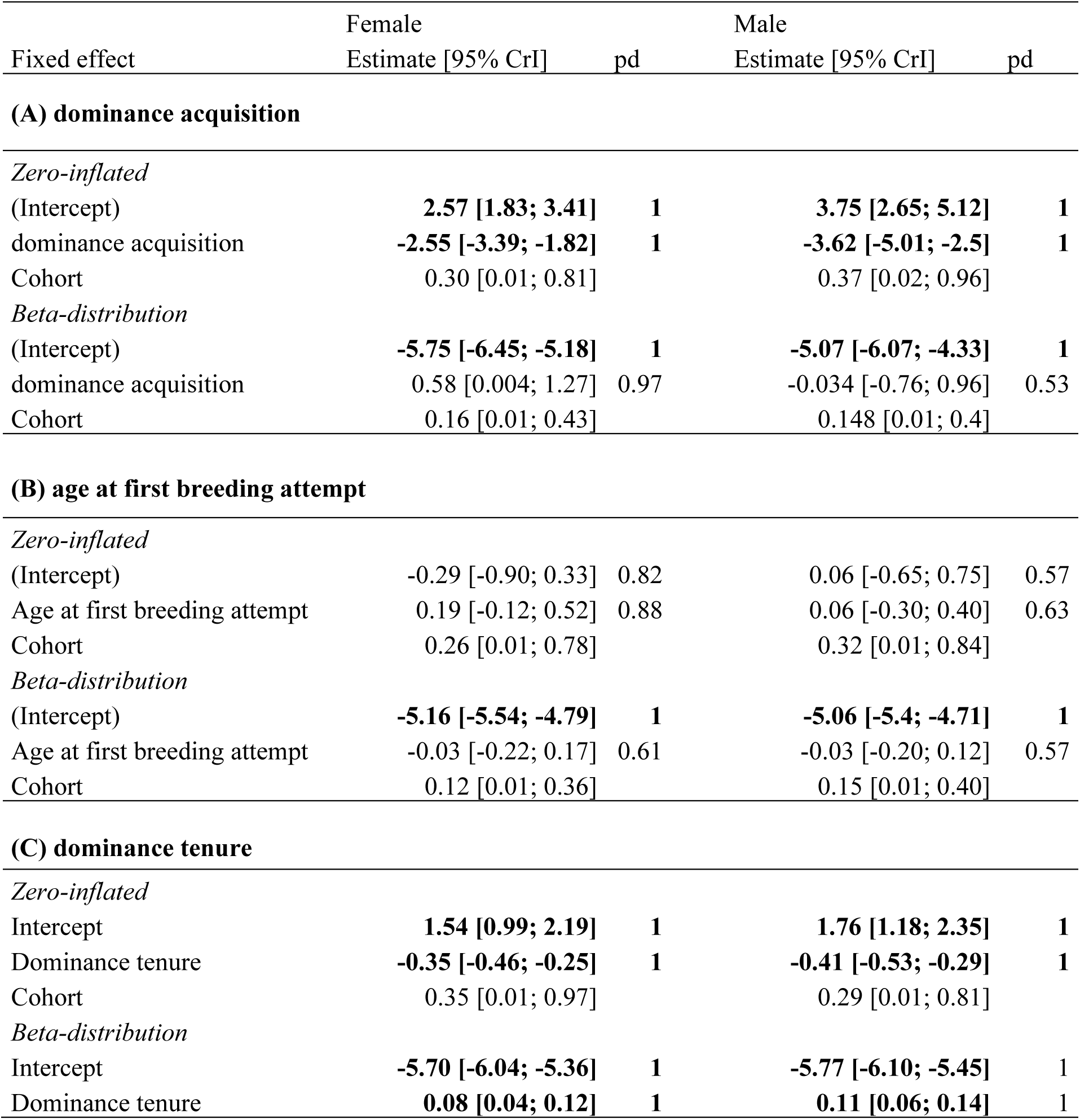
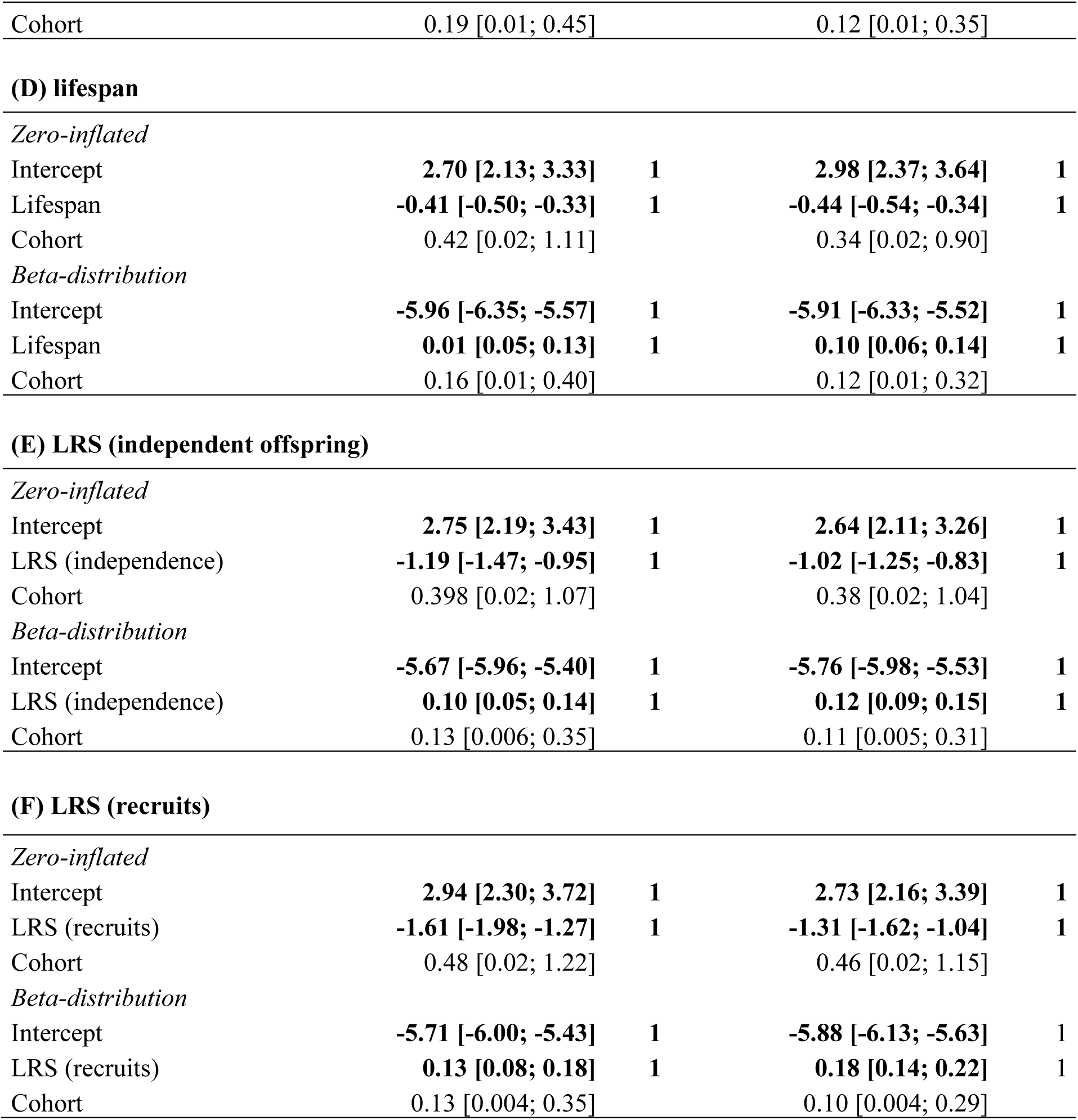
The relationship between (A) dominance acquisition, (B) age at first breeding attempt, (C) dominance tenure, (D) lifespan, (E) LRS (independent offspring ≥3 months old),and (F) LRS (recruits ≥1yr old), and the estimated individual genetic contribution (IGC) to the population 15 years (∼3 generations) in the future in a closed population of Seychelles warblers. The beta distribution of the model assesses how the life-history traits influence future IGC for individuals with non-zero IGC; the zero-inflated portion of the model assesses how these influence the probability of an individual having an IGC of zero (indicating lineage extinction). The estimate, 95% credible interval (CrI), and probability of direction (pd) adjusted for multiple testing are given. Reference level for dominance = No. For the random cohort effect, standard deviations (and 95% CrIs) are shown. Effects with pd > 0.975 are considered significant and are shown in bold.

| Fixed effect | Female<br>Estimate [95% CrI] | pd | Male<br>Estimate [95% CrI] | pd |
| --- | --- | --- | --- | --- |
| **(A) dominance acquisition** |  |  |  |  |
| *Zero-inflated* |  |  |  |  |
| (Intercept) | **2.57 [1.83; 3.41]** | **1** | **3.75 [2.65; 5.12]** | **1** |
| dominance acquisition | **-2.55 [-3.39; -1.82]** | **1** | **-3.62 [-5.01; -2.5]** | **1** |
| Cohort | 0.30 [0.01; 0.81] |  | 0.37 [0.02; 0.96] |  |
| *Beta-distribution* |  |  |  |  |
| (Intercept) | **-5.75 [-6.45; -5.18]** | **1** | **-5.07 [-6.07; -4.33]** | **1** |
| dominance acquisition | 0.58 [0.004; 1.27] | 0.97 | -0.034 [-0.76; 0.96] | 0.53 |
| Cohort | 0.16 [0.01; 0.43] |  | 0.148 [0.01; 0.4] |  |
| **(B) age at first breeding attempt** |  |  |  |  |
| *Zero-inflated* |  |  |  |  |
| (Intercept) | -0.29 [-0.90; 0.33] | 0.82 | 0.06 [-0.65; 0.75] | 0.57 |
| Age at first breeding attempt | 0.19 [-0.12; 0.52] | 0.88 | 0.06 [-0.30; 0.40] | 0.63 |
| Cohort | 0.26 [0.01; 0.78] |  | 0.32 [0.01; 0.84] |  |
| *Beta-distribution* |  |  |  |  |
| (Intercept) | **-5.16 [-5.54; -4.79]** | **1** | **-5.06 [-5.4; -4.71]** | **1** |
| Age at first breeding attempt | -0.03 [-0.22; 0.17] | 0.61 | -0.03 [-0.20; 0.12] | 0.57 |
| Cohort | 0.12 [0.01; 0.36] |  | 0.15 [0.01; 0.40] |  |
| **(C) dominance tenure** |  |  |  |  |
| *Zero-inflated* |  |  |  |  |
| Intercept | **1.54 [0.99; 2.19]** | **1** | **1.76 [1.18; 2.35]** | **1** |
| Dominance tenure | **-0.35 [-0.46; -0.25]** | **1** | **-0.41 [-0.53; -0.29]** | **1** |
| Cohort | 0.35 [0.01; 0.97] |  | 0.29 [0.01; 0.81] |  |
| *Beta-distribution* |  |  |  |  |
| Intercept | **-5.70 [-6.04; -5.36]** | **1** | **-5.77 [-6.10; -5.45]** | **1** |
| Dominance tenure | **0.08 [0.04; 0.12]** | **1** | **0.11 [0.06; 0.14]** | **1** |
| Cohort | 0.19 [0.01; 0.45] |  | 0.12 [0.01; 0.35] |  |
| **(D) lifespan** |  |  |  |  |
| *Zero-inflated* |  |  |  |  |
| Intercept | 2.70 [2.13; 3.33] | 1 | 2.98 [2.37; 3.64] | 1 |
| Lifespan | -0.41 [-0.50; -0.33] | 1 | -0.44 [-0.54; -0.34] | 1 |
| Cohort | 0.42 [0.02; 1.11] |  | 0.34 [0.02; 0.90] |  |
| *Beta-distribution* |  |  |  |  |
| Intercept | -5.96 [-6.35; -5.57] | 1 | -5.91 [-6.33; -5.52] | 1 |
| Lifespan | 0.01 [0.05; 0.13] | 1 | 0.10 [0.06; 0.14] | 1 |
| Cohort | 0.16 [0.01; 0.40] |  | 0.12 [0.01; 0.32] |  |
| **(E) LRS (independent offspring)** |  |  |  |  |
| *Zero-inflated* |  |  |  |  |
| Intercept | 2.75 [2.19; 3.43] | 1 | 2.64 [2.11; 3.26] | 1 |
| LRS (independence) | -1.19 [-1.47; -0.95] | 1 | -1.02 [-1.25; -0.83] | 1 |
| Cohort | 0.398 [0.02; 1.07] |  | 0.38 [0.02; 1.04] |  |
| *Beta-distribution* |  |  |  |  |
| Intercept | -5.67 [-5.96; -5.40] | 1 | -5.76 [-5.98; -5.53] | 1 |
| LRS (independence) | 0.10 [0.05; 0.14] | 1 | 0.12 [0.09; 0.15] | 1 |
| Cohort | 0.13 [0.006; 0.35] |  | 0.11 [0.005; 0.31] |  |
| **(F) LRS (recruits)** |  |  |  |  |
| *Zero-inflated* |  |  |  |  |
| Intercept | 2.94 [2.30; 3.72] | 1 | 2.73 [2.16; 3.39] | 1 |
| LRS (recruits) | -1.61 [-1.98; -1.27] | 1 | -1.31 [-1.62; -1.04] | 1 |
| Cohort | 0.48 [0.02; 1.22] |  | 0.46 [0.02; 1.15] |  |
| *Beta-distribution* |  |  |  |  |
| Intercept | -5.71 [-6.00; -5.43] | 1 | -5.88 [-6.13; -5.63] | 1 |
| LRS (recruits) | 0.13 [0.08; 0.18] | 1 | 0.18 [0.14; 0.22] | 1 |
| Cohort | 0.13 [0.004; 0.35] |  | 0.10 [0.004; 0.29] |  |

We then tested whether some life-history traits better predicted IGC after 15 years and whether this differed between sexes. For both sexes, age at first breeding attempt was the poorest predictor of IGC (R^2^ for both sexes = 2%, Table 3), with acquiring dominance better predicting IGC than age at first breeding attempt (R^2^ for both sexes = 11%, Figure 3A and B; Table 3). In turn, both dominance tenure and lifespan were better predictors of IGC than acquiring dominance but did not significantly differ in their predictive power (lifespan: males, 37.3%, females, 37.4%; dominant breeding tenure: males, 34.1%, females, 27.5%, Figure 3C, D; Table 3). Furthermore, for only males, both LRS measures better predicted IGC than all other proxies [LRS (independent offspring) = 61%, LRS (recruits) = 62%] and the predictive power of both LRS traits were higher in males than in females (females: LRS (independent offspring) = 46%, LRS (recruits) = 49%, Figure 3E and F; Table 3; Table 4). Across both sexes there was no difference in the extent to which LRS calculated by censusing either independents or recruits predicted IGC (Figure 3E and F; Table 3).

**Figure 3.**
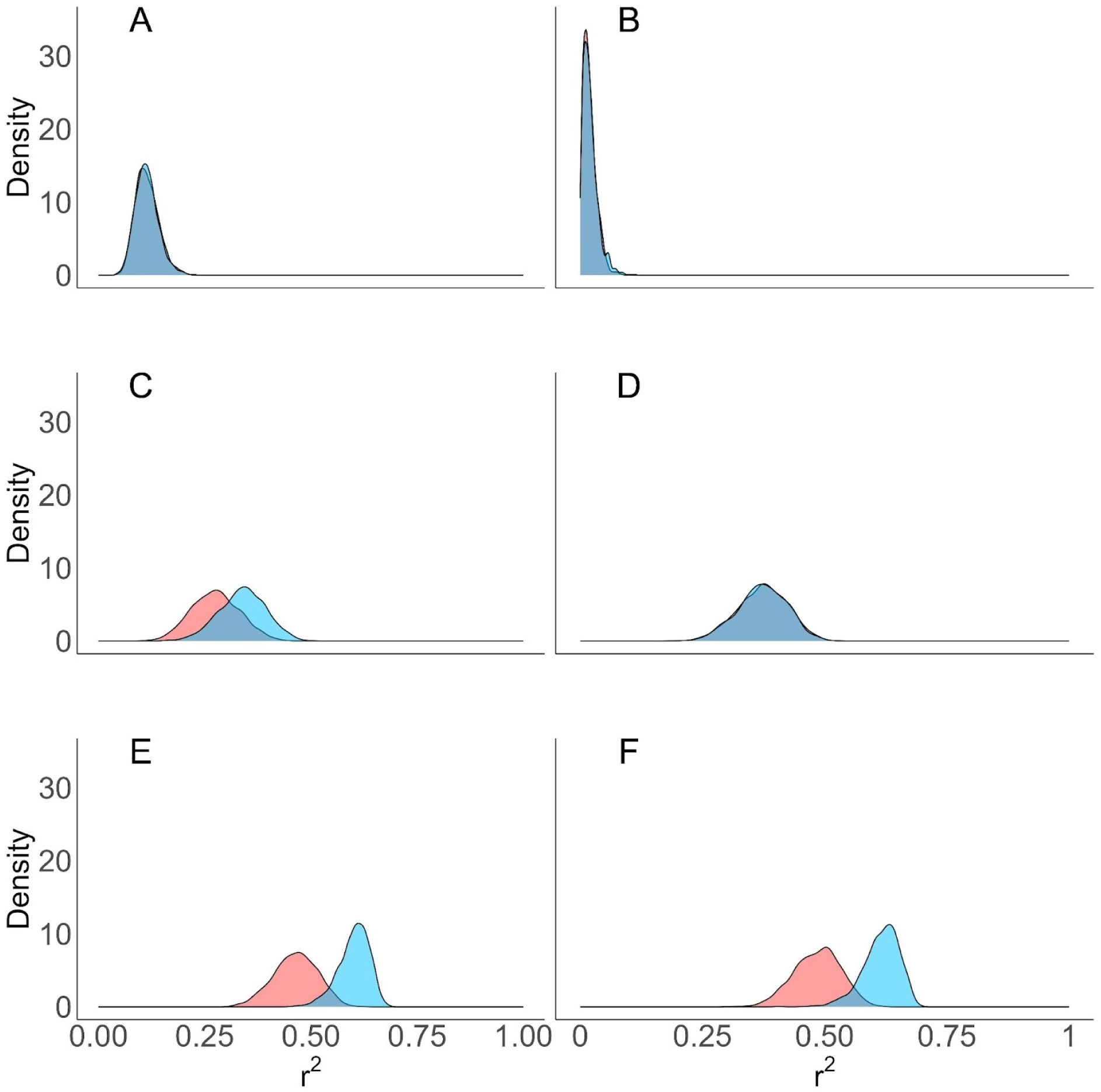
The density distributions of the R^2^ for individual genetic contributions (IGC) predicted 15 years in the future, for female (red) and male (blue) Seychelles warblers, by (A) dominance acquisition, (B) age at first breeding attempt, (C) dominance tenure, (D) lifespan, (E) LRS (independent offspring ≥3 months old; and (F) LRS (recruits ≥1 year old).

**Table 3:** Results from zero-inflated beta distribution models for female and male Seychelles warblers assessing the ability of life-history traits (A-F) to predict individual genetic contributions (IGC) 15 years in the future, and a comparison of the predictive ability of the different life-history traits. On the diagonal are the R^2^ values from the model with the life-history trait. Below the diagonal are the pairwise comparisons of R^2^ between models using different life-history traits (ΔR^2^). Ninety-five per cent credible intervals are shown in square brackets; bold values indicate significance. LRS = lifetime reproductive success: independent = offspring: 3 months or older; recruits = offspring: 1 year or older.

|  | (A) Dominance acquisition | (B) Age at first breeding attempt | (C) Dominance tenure | (D) Lifespan | (E) LRS (independent offspring) | (F) LRS (recruits) |
| --- | --- | --- | --- | --- | --- | --- |
| **Females** |  |  |  |  |  |  |
| (A) Dominance acquisition | 0.11 [0.07; - 0.18] |  |  |  |  |  |
| (B) Age at first breeding attempt | **0.10 [0.04; 0.16]** | 0.016 [0.00; 0.05] |  |  |  |  |
| (C) Dominance tenure | **-0.16 [-0.28; -0.04]** | **-0.26 [-0.37; -0.15]** | 0.28 [0.17; 0.39] |  |  |  |
| (D) Lifespan | **-0.26 [-0.37; -0.14]** | **-0.35 [-0.46; -0.25]** | -0.1 [-0.25; 0.06] | 0.37 [0.27; 0.47] |  |  |
| (E) LRS (independent offspring) | **-0.35 [-0.45; -0.24]** | **-0.44 [-0.54; -0.34]** | **-0.19 [-0.33; -0.04]** | -0.09 [-0.23; 0.05] | 0.47 [0.36; 0.56] |  |
| (F) LRS (recruits) | **-0.37 [-0.47; -0.26]** | **-0.47 [-0.55; -0.37]** | **-0.21 [-0.34; -0.06]** | -0.11 [-0.25; 0.03] | -0.02 [-0.15; 0.12] | 0.49 [0.39; 0.57] |
| **Males** |  |  |  |  |  |  |
| (A) Dominance acquisition | 0.112 [0.07; 0.17] |  |  |  |  |  |
| (B) Age at first breeding attempt | **0.09 [0.03; 0.16]** | 0.018 [0.00; 0.06] |  |  |  |  |
| (C) Dominance tenure | **-0.23 [-0.34; -0.1]** | **-0.32 [-0.42; -0.2]** | 0.341 [0.23; 0.44] |  |  |  |
| (D) Lifespan | **-0.26 [-0.37; -0.15]** | **-0.35 [-0.45; -0.24]** | -0.03 [-0.17; 0.11] | 0.373 [0.27; 0.47] |  |  |
| (E) LRS (independent offspring) | **-0.49 [-0.57; -0.39]** | **-0.58 [-0.64; -0.49]** | **-0.26 [-0.39; -0.13]** | **-0.23 [-0.34; -0.11]** | 0.606 [0.52; 0.66] |  |
| (F) LRS (recruits) | **-0.50 [-0.58; -0.4]** | **-0.59 [-0.66; -0.5]** | **-0.28 [-0.4; -0.15]** | **-0.24 [-0.36; -0.12]** | -0.01 [-0.12; 0.09] | 0.62 [0.53; 0.67] |
| Sex-ΔR2 | 0.001 [-0.07; 0.08] | <sup>^</sup>-0.002 [-0.04; 0.04] | -0.063 [-0.22; 0.09] | 0.003 [-0.13; 0.14] | **-0.14 [-0.26; -0.02]** | **-0.13 [-0.24; -0.01]** |

## DISCUSSION

We used complete life-history records and genetic pedigree data spanning 26 years to quantify the genetic contributions (IGC) after 15 years across 11 birth cohorts in a cooperatively breeding bird (Seychelles warbler) (N = 680). Across both sexes, we then quantified to what extent these genetic contributions were predicted by six life-history traits: (A) acquisition of a dominant breeding position, (B) age at first breeding attempt, (C) tenure as a dominant breeder, (D) lifespan, and (E) LRS (measured as both the number of independent [surviving 3 months+] and (F) recruited [surviving 1 year+] offspring produced over an individual’s lifetime, Table 1). Across both sexes, we found that almost all life-history traits predicted both the probability of an individual’s genetic lineage going extinct (except for age at first breeding attempt, Figure 1), and the amount of IGC in those individuals whose lineages did not go extinct (except for age at first breeding attempt and dominance acquisition, Figure 2). However, life-history traits varied considerably in their ability to predict IGC (2-62%), with both measures of lifetime reproductive success also better predicting IGC in males than females (Figure 3). Together these results give insight into both the individual and population-level processes shaping future gene pools.

As expected, LRS was a strong predictor of IGC in the Seychelles warbler suggesting that natural selection plays an important role in long-term evolutionary processes within this population. Moreover, at least for males, the amount of variation in IGC explained by LRS in the Seychelles warbler was higher than in previous studies on other species. For example, in the Seychelles warbler, LRS (recruited offspring) explained ∼49% of the variation in IGC for females and 62% for males, whereas for the Mandarte song sparrows (*Melospiza melodia*) this was ∼48% for females and 55% for males, even though both studies measured recruits as the same: offspring surviving for more than one year (Reid et al., 2019). One methodological explanation for this difference is that IGC for the song sparrows were calculated over a longer period (20 versus 15 years) and more generations (8 versus 3), potentially increasing the role of stochastic and demographic forces to dissociate LRS from IGC. Indeed, LRS explained only 28-32% of the variation in IGC when measured after ∼10 generations in humans (Young et al., 2023) but 52-62% when measured after two generations in Bighorn sheep [*Ovis canadensis*, (van de Walle et al., 2022)].

Biological differences between populations may also play a role. LRS should be a better predictor of IGC in stable populations (Brommer et al., 2002; Mcgraw & Caswell, 1996; van de Walle et al., 2022), and the Seychelles warbler population on Cousin has, indeed, remained largely stable and at carrying capacity over the length of the study period (see *Methods* or Komdeur et al., 2016). Whilst LRS (recruited offspring) was a strong predictor of IGC within our study, this finding is not always echoed by studies considering systems with unstable population sizes. For instance, in Lundy house sparrows (*Passer domesticus*), LRS (recruited offspring) explained only 21% of the variation in IGC, which is likely due to one season of raptor predation lowering the survival of the descendants of some focal individuals and thereby lowering their IGC independent of their reproductive success (Alif et al., 2022). Another important factor is migration, where the ability of life-history traits to predict long-term genetic contributions may reduce if high emigration or immigration disrupt signals of natural selection from the ancestral population. Indeed, migration is virtually non-existent in the Seychelles warbler, while being high in the Mandarte song sparrow population (Reid et al., 2021). However, whilst LRS (recruited offspring) was among the life-history traits that best predicted IGC within our study, there was still substantial unexplained variation in IGC (∼38% for males and 51% for females). This may in part be due to the week correlation between the LRS of individuals and their descendants (Reid et al., 2019), with one recent comparative study suggesting that, on average, only ∼3% of variance in LRS is due to genetic variation in wild animal populations (Bonnet et al., 2022). Therefore, future studies should aim to go beyond examining how well life-history traits (or “fitness proxies”) predict IGC, and instead focus on the role of stochasticity (e.g., genetic drift) and the environment, which has largely been overlooked (but see (Borger et al., 2023)).

The ability of LRS to predict IGC also varied between sexes: LRS (independent and recruited offspring) explained ∼61% and 62% of the variation in IGC for males, but only ∼47% and 49% for females, respectively. For males, the predictive power of both LRS measures was also higher than for lifespan, but for females, it was not. A likely explanation for this is that females usually only have one clutch per breeding season, which is usually limited to one offspring (Komdeur, 1994), meaning that LRS for females is largely determined by their lifespan. Some males, however, can gain higher LRS through extra-pair paternity or lower when cuckolded (Raj Pant et al., 2022), increasing the variance in annual reproductive success amongst males and meaning that both LRS measures better predicted IGC than lifespan in males. This demonstrates how sex differences can impact fundamental parameters in evolutionary biology, such as variance in fitness and future contributions to gene pools.

Although lifespan and dominance were worse predictors of IGC than LRS in males, they were better predictors of IGC than age at first breeding attempt across both sexes, suggesting that the duration of life and of dominant breeding were far more important than the age of starting breeding in constraining LRS and subsequent IGC. In fact, age at first breeding attempt did not affect either the probability of lineage extinction nor the IGC of surviving lineages (Table 2). One possible reason for this is a trade-off between early and late-life fitness components (Lemaître et al., 2015; Stearns, 1992; Williams, 1966), which would mean that individuals reproducing earlier in life suffer reduced lifespans or reproductive success in later life. Indeed, in the Seychelles warbler, an earlier age at first breeding is negatively associated with both late-life survival (Hammers et al., 2013) and male LRS (Raj Pant et al., 2019). If true, this highlights the importance of considering life-history trade-offs when using life-history traits to estimate selection.

Finally, we draw the attention of readers to some limitations in our study. First, we calculated IGC as those that contributed to direct descent. However, it is important to acknowledge that individuals in our population may also gain indirect genetic contributions through helping relatives (Hamilton, 1964; Richardson et al., 2003) and that the most important life-history traits in predicting IGC may differ when considering indirect genetic contributions. Second, while we found no differences in predictive power between LRS when censusing independent or recruited offspring, we believe this represents the limitations of how early we are able to measure LRS, rather than timing of censusing not being important *per se*. In our system, we can only reliably quantify the number of offspring in nests after three months, and censusing LRS at earlier stages would have resulted in biased estimates. Indeed, other studies have shown that the age of censusing offspring LRS does have implications for how well LRS predicts IGC (Alif et al., 2022; Reid et al., 2019). Therefore, we urge researchers to both carefully consider which LRS measure is most reliable for estimating selection in their system - in the light of previously highlighted issues (Hadfield, 2008; Wolf & Wade, 2001) - and clearly specify when exactly they have censused offspring for calculating LRS.

In summary, this study has extensively quantified how well different life histories predict long-term genetic contributions in a cooperatively breeding system. This paper highlights the diversity of journeys that wild animals - and especially cooperative breeders - take from birth to death and helps to elucidate how different parts of this journey ultimately affect long-term evolutionary processes. This adds to the evidence of the importance of long-term individual-level data on wild organisms for studying evolution (Sheldon et al., 2022). However, despite using extensive and detailed long-term pedigree data from a wild population, our study is still limited to showing the relationships between life-history traits and IGC only over a few generations. It is therefore vital to continue to extend these data into future years, which will be key to better refining our understanding of evolution over evolutionary scales.

## Author contributions

The project was conceived by EC and HLD, with input from AMS. The long-term field work was managed by DSR, with DSR, EC, TB, and HLD contributing to data collection. Molecular genotyping was undertaken by MV and DSR. Parentage analyses were run by HLD and SFW. Statistical analyses were designed and run by EC, EAY, MAV, and HLD with input from AMS and DSR. EC wrote the first draft, which was then rewritten by EAY, MAV, and HLD and commented on by EC, MV, MAV, SFW, TB, JK, and DSR. All authors agreed to submission.

## Acknowledgements

We thank all the researchers who collected the long-term data, Ian Stevenson and Owen Howison for maintaining the database, and Darren Hunter for providing R code for estimating IGC. Susan Johnston for editing genedroppeR for use with multiple breeding seasons per year, and Alex Sparks for discussion and analytical insights. Ben Hatchwell and Elizabeth Duncan who provided feedback on an earlier draft. Thanks to Nature Seychelles for providing access to the Cousin Island Nature Reserve, and the Seychelles Department of Environment and the Seychelles Bureau of Standards for approving the protocol locally and providing research permits. E.A.Y.’s PhD was funded by the University of Groningen, through a Rosalind Franklin Fellowship awarded to H.L.D. EC was supported by a Leeds Doctoral Scholarship and a Climate Research Bursary Fund, University of Leeds. The long-term data gathering that enabled this study was supported by various NERC grants including NE/B504106/1 (TB and DSR), NE/I021748/1 (HLD), and NE/F02083X/1 and NE/K005502/1 (DSR); as well as a NWO Rubicon (825.09.013), NWO visitors grant (040.11.232 to JK and HLD), and NWO grants (854.11.003 and 823.01.014 to JK).

## REFERENCES

Alif, Ž., Dunning, J., Chik, H. Y. J., Burke, T., & Schroeder, J. (2022). What is the best fitness measure in wild populations? A case study on the power of short-term fitness proxies to predict reproductive value. PLoS One, 17(4), e0260905. 10.1371/journal.pone.0260905

Bonnet, T., Morrissey, M. B., Villemereuil, P. D., Alberts, S. C., Arcese, P., Bailey, L. D., Boutin, S., Brekke, P., Brent, L. J. N., Camenisch, G., Charmantier, A., Clutton-Brock, T. H., Cockburn, A., Coltman, D. W., Courtiol, A., Davidian, E., Evans, S. R., Ewen, J. G., Festa-bianchet, M., … Kruuk, L. E. B. (2022). Genetic variance in fitness indicates rapid on-going adaptive evolution in wild animals. Science, 376(6596), 1012–1016.

Brommer, J. E., Gustafsson, L., Pietiäinen, H., & Merilä, J. (2004). Single-generation estimates of individual fitness as proxies for long-term genetic contribution. American Naturalist, 163(4), 505–517. 10.1086/382547

Brouwer, L., Barr, I., Van De Pol, M., Burke, T., Komdeur, J., & Richardson, D. S. (2010). MHC-dependent survival in a wild population: Evidence for hidden genetic benefits gained through extra-pair fertilizations. Molecular Ecology, 19(16), 3444–3455.

Brouwer, L., Richardson, D. S., Eikenaar, C., & Komdeur, J. (2006). The role of group size and environmental factors on survival in a cooperatively breeding tropical passerine. Journal of Animal Ecology, 75(6), 1321–1329. 10.1111/j.1365-2656.2006.01155.x

Bürkner, P. C. (2018). Advanced Bayesian multilevel modeling with the R package brms. R Journal, 10(1), 395–411. 10.32614/rj-2018-017

Chen, N., Juric, I., Cosgrove, E. J., Bowman, R., Fitzpatrick, J. W., Schoech, S. J., Clark, A. G., & Coop, G. (2019). Allele frequency dynamics in a pedigreed natural population. Proceedings of the National Academy of Sciences of the United States of America, 116(6), 2158–2164. 10.1073/pnas.1813852116

Chesterton, E., Sparks, A. M., Burke, T., Komdeur, J., Richardson, D. S., & Dugdale, H. L. (2024). The impact of helping experience on helper life-history and fitness in a cooperatively breeding bird. Evolution, 78(4), 690–700. 10.1093/evolut/qpad199

Clutton-Brock, T. H. (1988). Reproductive success: Studies of individual variation in contrasting breeding systems. University of Chicago Press.

Crow, J. (1958). Index of total selection intensity-some possibilities for measuring selection intensities in man. Human Biology, 30, 1–3.

Davies, C. S., Taylor, M. I., Hammers, M., Burke, T., Komdeur, J., Dugdale, H. L., & Richardson, D. S. (2021). Contemporary evolution of the innate immune receptor gene TLR3 in an isolated vertebrate population. Molecular Ecology, 30(11), 2528–2542. 10.1111/mec.15914

Dugdale, H. L., Pope, L. C., Newman, C., MacDonald, D. W., & Burke, T. (2011). Age-specific breeding success in a wild mammalian population: Selection, constraint, restraint and senescence. Molecular Ecology, 20(15), 3261–3274. 10.1111/j.1365-294X.2011.05167.x

Evans, S. R., & Postma, E. (2024). Counting chicks before they hatch: Extending the observed lifetime to better characterize evolutionary processes in the wild. Evolution, 79(2), 155–163. 10.1093/evolut/qpae171

Gates, J. E., & Gysel, L. W. (1978). Avian nest dispersion and fledging success in field-forest ecotones. Ecology, 59(5), 871–883.

Gelman, A., Goodrich, B., Gabry, J., & Vehtari, A. (2019). R-squared for Bayesian Regression Models. The American Statistician, 73(3), 307–309. 10.1080/00031305.2018.1549100

Grafen, A. (1988). On the uses of data on lifetime reproductive success. In T. Clutton-brock (Ed.), Reproductive success (University of Chicago Press, Chicago, IL, pp. 454–751).

Grant, B. R. (1985). Selection on bill characters in a population of Darwin’s finches: Geospiza conirostris on Isla Genovesa, Galapagos. Evolution, 39(3), 523–532.

Hadfield, J. D. (2008). Estimating evolutionary parameters when viability selection is operating. Proceedings of the Royal Society B: Biological Sciences, 275(1635), 723–734. 10.1098/rspb.2007.1013

Hadfield, J. D., Richardson, D., & Burke, T. (2006). Towards unbiased parentage assignment: Combining genetic, behavioural and spatial data in a Bayesian framework. Molecular Ecology, 15(12), 3715–3730.

Hamilton, W. D. (1964). The genetical evolution of social behaviour. II. Journal of Theoretical Biology, 7(1), 17–52. 10.1016/0022-5193(64)90039-6

Hammers, M., Richardson, D. S., Burke, T., & Komdeur, J. (2013). The impact of reproductive investment and early-life environmental conditions on senescence: Support for the disposable soma hypothesis. Journal of Evolutionary Biology, 26(9), 1999–2007. 10.1111/jeb.12204

Hawkes, K., O’Connell, J. F., Jones, N. G. B., Alvarez, H., & Charnov, E. L. (1998). Grandmothering, menopause, and the evolution of human life histories. Proceedings of the National Academy of Sciences, 95(3), 1336–1339. 10.1073/pnas.95.3.1336

Hudson, R. R. (1990). Gene genealogies and the coalescent process. Oxford Surveys in Evolutionary Biology, 7(1), 44.

Hunter, D. C., Pemberton, J. M., Pilkington, J. G., & Morrissey, M. B. (2019). Pedigree-Based Estimation of Reproductive Value. Journal of Heredity, 110(4), 433–444. 10.1093/jhered/esz033

Johnston, S. E. (2020). genedroppeR: An R package to run single-locus genedrop analyses through complex pedigrees (Version 0.1.0) [Computer software]. https://github.com/susjoh/genedroppeR

Kingsolver, J. G., Hoekstra, H. E., Hoekstra, J. M., Berrigan, D., Vignieri, S. N., Hill, C. E., Hoang, A., Gibert, P., & Beerli, P. (2001). The strength of phenotypic selection in natural populations. American Naturalist, 157(3), 245–261. 10.1086/319193

Komdeur, J. (1992). Importance of habitat saturation and territory quality for evolution of cooperative breeding in the Seychelles warbler. Nature, 358(6386), 493–495.

Komdeur, J. (1994). Conserving the Seychelles warbler Acrocephalus sechellensis by translocation from Cousin Island to the islands of Aride and Cousine. Biological Conservation, 67(2), 143–152. 10.1016/0006-3207(94)90360-3

Komdeur, J. (2003). Adaptations and maladaptations to island living in the Seychelles Warbler. Ornithological Science, 2(2), 79–88. 10.2326/osj.2.79

Komdeur, J., Burke, T., Dugdale, H., & Richardson, D. S. (2016). Seychelles warblers: Complexities of the helping paradox. In W. D. Koenig & J. L. Dickinson (Eds), Cooperative Breeding in Vertebrates (1st edn, pp. 197–216). Cambridge University Press. 10.1017/cbo9781107338357.013

Komdeur, J., Piersma, T., Kraaijeveld, K., Kraaijeveld-Smit, F., & Richardson, D. S. (2004). Why Seychelles Warblers fail to recolonize nearby islands: Unwilling or unable to fly there? Ibis, 146(2), 298–302.

Lack, D. (1947). The significance of clutch-size. Ibis, 89(2), 302–352. 10.1111/j.1474-919X.1947.tb04155.x

Lande, R., & Arnold, S. J. (1983). The measurement of selection on correlated characters. Evolution, 37(6), 1201226. https://www.jstor.org/stable/2408842

Lansing, A. I. (1947). A transmissible, cumulative, and reversible factor in aging. Journal of Gerontology, 2(3), 228–239. 10.1093/geronj/2.3.228

Lemaître, J. F., Berger, V., Bonenfant, C., Douhard, M., Gamelon, M., Plard, F., & Gaillard, J. M. (2015). Early-late life trade-offs and the evolution of ageing in the wild. Proceedings of the Royal Society B: Biological Sciences, 282(1806). 10.1098/rspb.2015.0209

Makowski, D., Ben-Shachar, M. S., Chen, S. H. A., & Lüdecke, D. (2019). Indices of effect existence and significance in the bayesian framework. Frontiers in Psychology, 10(December), 1–14. 10.3389/fpsyg.2019.02767

Meierhofer, I., Schwarz, H. H., & Müller, J. K. (1999). Seasonal variation in parental care, offspring development, and reproductive success in the burying beetle, Nicrophorus vespillo. Ecological Entomology, 24(1), 73–79.

Monaghan, P., Maklakov, A. A., & Metcalfe, N. B. (2020). Intergenerational transfer of ageing: Parental age and offspring lifespan. Trends in Ecology & Evolution, 35(10), 927–937. 10.1016/j.tree.2020.07.005

Mumme, RonaldL. (1992). Do helpers increase reproductive success?: An experimental analysis in the Florida scrub jay. Behavioral Ecology and Sociobiology, 31(5). 10.1007/bf00177772

R Core Team. (2020). R: A language environment for statistical computing [Computer software]. R Foundation for Statistical Computing. http://www.r-project.org/.

Raj Pant, S., Komdeur, J., Burke, T. A., Dugdale, H. L., & Richardson, D. S. (2019). Socio-ecological conditions and female infidelity in the Seychelles warbler. Behavioral Ecology, 30(5), 1254–1264.

Raj Pant, S., Versteegh, M. A., Hammers, M., Burke, T., Dugdale, H. L., Richardson, D. S., & Komdeur, J. (2022). The contribution of extra-pair paternity to the variation in lifetime and age-specific male reproductive success in a socially monogamous species. Evolution, 76(5), 915–930. 10.1111/evo.14473

Reid, J. M., Arcese, P., Nietlisbach, P., Wolak, M. E., Muff, S., Dickel, L., & Keller, L. F. (2021). Immigration counter-acts local micro-evolution of a major fitness component: Migration-selection balance in free-living song sparrows. Evolution Letters, 5(1), 48–60. 10.1002/evl3.214

Reid, J. M., Nietlisbach, P., Wolak, M. E., Keller, L. F., & Arcese, P. (2019). Individuals’ expected genetic contributions to future generations, reproductive value, and short-term metrics of fitness in free-living song sparrows (Melospiza melodia). Evolution Letters, 3(3), 271–285. 10.1002/evl3.118

Richardson, D. S., Burke, T., & Komdeur, J. (2002). Direct benefits and the evolution of female-biased cooperative breeding in Seychelles warblers. Evolution, 56(11), 2313–2321.

Richardson, D. S., Jury, F., Blaakmeer, K., Komdeur, J., & Burke, T. (2001). Parentage assignment and extra-group paternity in a cooperative breeder: The Seychelles warbler (Acrocephalus sechellensis). Molecular Ecology, 10(9), 2263–2273.

Safina, C., & Burger, J. (1983). Effects of human disturbance on reproductive success in the black skimmer. The Condor, 85(2), 164–171. 10.2307/1367250

Sheldon, B. C., Kruuk, L. E. B., & Alberts, S. C. (2022). The expanding value of long-term studies of individuals in the wild. Nature Ecology & Evolution, 6(12), 1799–1801. 10.1038/s41559-022-01940-7

Sparks, A. M., Hammers, M., Komdeur, J., Burke, T., Richardson, D. S., & Dugdale, H. L. (2022). Sex-dependent effects of parental age on offspring fitness in a cooperatively breeding bird. Evolution Letters, 6(6), 438–449. 10.1002/evl3.300

Sparks, A. M., Spurgin, L. G., van der Velde, M., Fairfield, E. A., Komdeur, J., Burke, T., Richardson, D. S., & Dugdale, H. L. (2022). Telomere heritability and parental age at conception effects in a wild avian population. Molecular Ecology, 31(23), 6324–6338.

Stan Development Team. (2020). RStan: The R interface to Stan [Computer software]. https://mc-stan.org/

Stearns, S. C. (1989). Trade-offs in life-history evolution. Functional Ecology, 3(3), 259. 10.2307/2389364

Stearns, S. C. (1992). The Evolution of Life Histories. Oxford University Press; WorldCat.

van de Walle, J., Larue, B., Pigeon, G., & Pelletier, F. (2022). Different proxies, different stories? Imperfect correlations and different determinants of fitness in bighorn sheep. Ecology and Evolution, 12(12), 1–12. 10.1002/ece3.9582

Williams, G. C. (1966). Natural selection, the costs of reproduction, and a refinement of Lack’s principle. American Naturalist, 100(916), 687–690. 10.2307/2459305

Wolf, J. B., & Wade, M. J. (2001). On the assignment of fitness to parents and offspring: Whose fitness is it and when does it matter? Journal of Evolutionary Biology, 14(2), 347–356. 10.1046/j.1420-9101.2001.00277.x

Young, E. A., Chesterton, E., Lummaa, V., Postma, E., & Dugdale, H. L. (2023). The long-lasting legacy of reproduction: Lifetime reproductive success shapes expected genetic contributions of humans after 10 generations. Proceedings of the Royal Society B: Biological Sciences, 290(1998), 20230287. 10.1098/rspb.2023.0287

