## Supplementary material for "Life-history traits as predictors of expected genetic contributions 15 years later in a cooperatively breeding bird"

### SUPPLEMENTARY MATERIALS

#### Supplementary methods: Gene-dropping

Gene-drop analyses were used to confirm that the microsatellites used to determine parentage were under neutral selection. The probability of a particular descendant having the same allele as the focal ancestor was estimated based on Mendelian principles. An offspring inherits one chromosome from each parent; if a heterozygous ancestor produces an offspring, there is a 50% chance of their offspring inheriting a particular autosomal allelic variant from them. From this, we can calculate the probability of a particular descendant within a lineage having an allelic variant, based on the variants of their ancestors. We used microsatellite data to test whether allelic variants were inherited at the rate we would expect if inheritance was random (i.e., the locus is under neutral selection). If a locus is not under neutral selection and confers a fitness advantage, we would expect the allelic variant to be present in the future population at a higher frequency. In contrast, if a variant confers a fitness disadvantage, we expect a reduced representation of the allelic variant in future generations. To test whether a particular locus was under neutral selection, gene-drop analyses were performed using the *R* package gendroppeR 0.1.0 [(Johnston, 2020) code available at https://github.com/susjoh/genedroppeR] on each of the 30 microsatellite loci used to determine parentage in the Seychelles warbler. All individuals within a pedigree (*n* = 1,853, cohorts: 1992–2018; Sparks, Spurgin, et al., 2022) were included within the analyses. We ran 1,000 simulations for each locus to estimate the confidence intervals. For all loci considered, the frequency of each allelic variant fell within the 95% confidence interval of what we would expect given random inheritance, indicating that all 30 loci are under neutral selection.

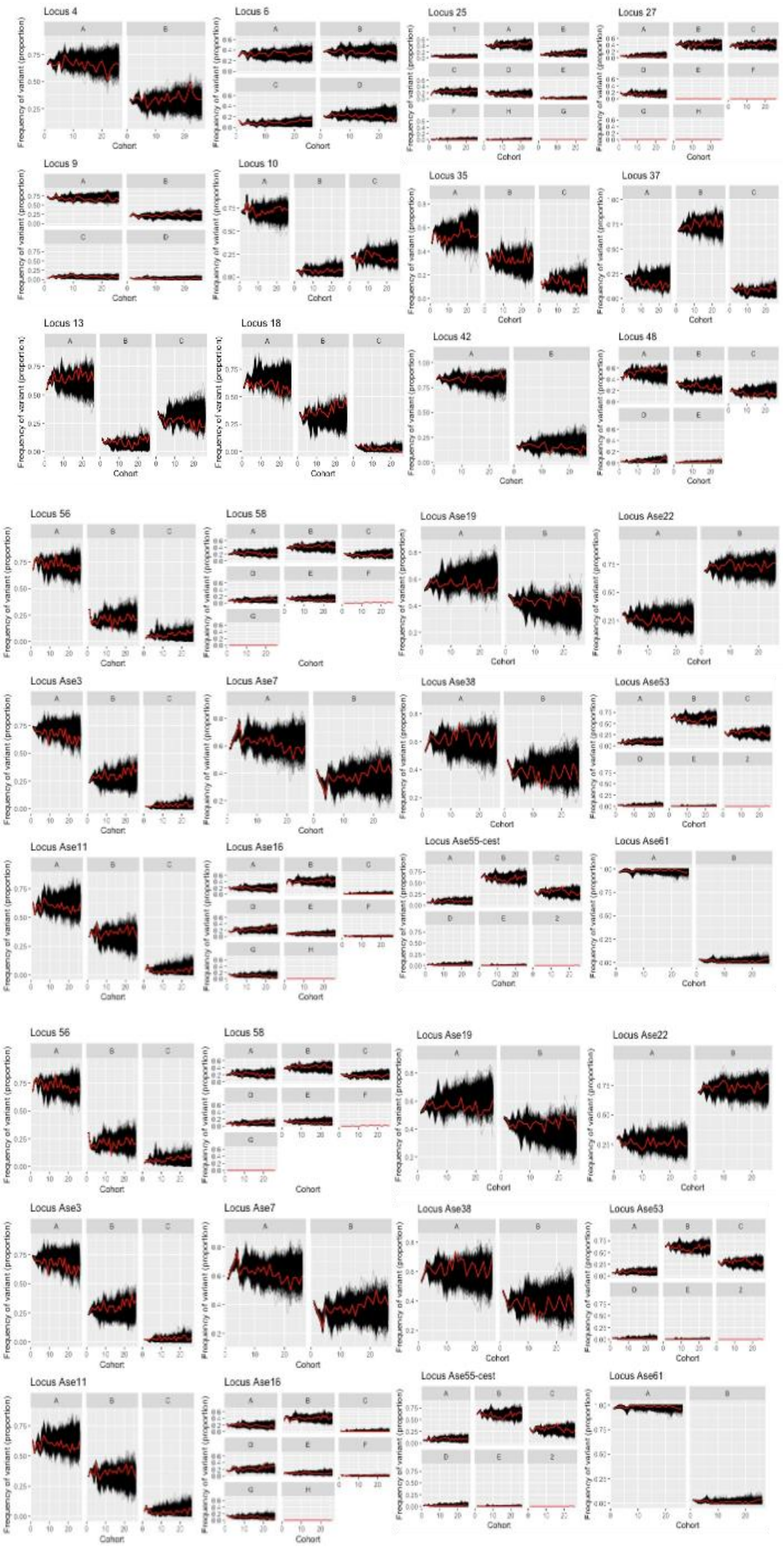


**Figure S1.** Results from gene-drop analysis for each microsatellite used to determine genetic parentage in the Seychelles warbler, to confirm that each allele is under neutral selection using methodology detailed in **Supplementary methods: Gene-dropping**.

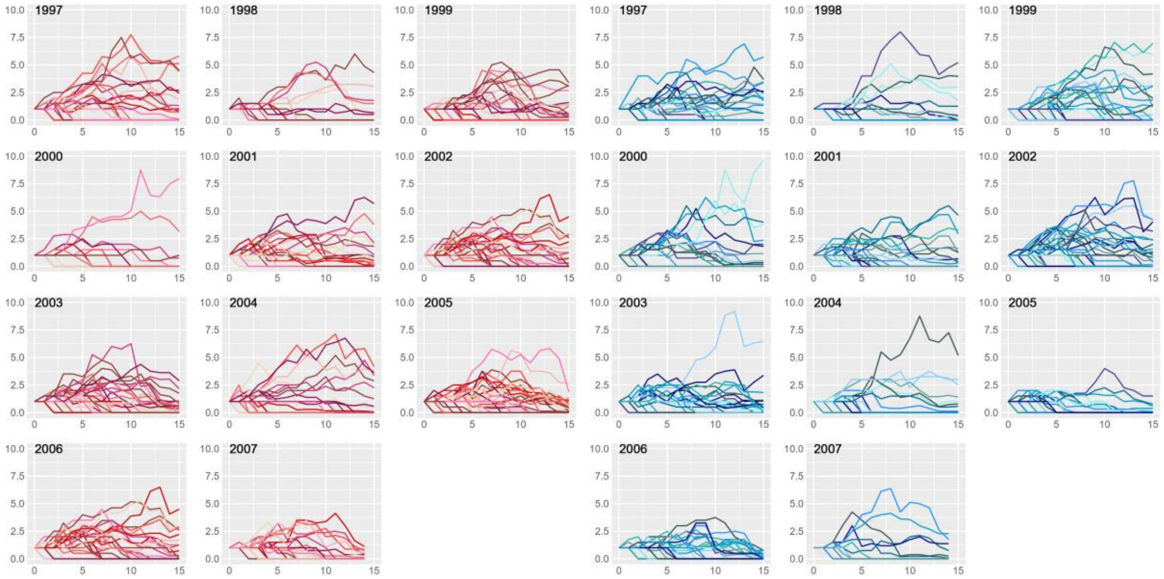


**Figure S2.** Change in absolute individual genetic contributions (IGC) over time in female (pink) and male (blue) Seychelles warblers over a 15-year timeframe for 11 cohorts (1997-2007), using methodology detailed in Hunter (2019).
